# Rapid aggregation of *Staphylococcus aureus* in synovial fluid is influenced by synovial fluid concentration, viscosity, and fluid dynamics-with evidence of polymer bridging

**DOI:** 10.1101/2022.02.02.478923

**Authors:** Amelia Staats, Peter W. Burback, Andrew Schwieters, Daniel Li, Anne Sullivan, Alexander R. Horswill, Paul Stoodley

## Abstract

Early bacterial survival in the post-surgical joint is still a mystery. Recently, synovial fluid-induced aggregation was proposed as a potential mechanism of bacterial protection upon entry into the joint. As synovial fluid is secreted back into the joint cavity following surgery, rapid fluctuations in synovial fluid concentration, composition, and viscosity occur. These changes, along with fluid movement from post-operative joint motion, will modify the environment and potentially affect the kinetics of aggregate formation. Through this work, we sought to evaluate the influence of exposure time, synovial fluid concentration, viscosity, and fluid dynamics on aggregation. Furthermore, we aimed to elucidate the primary mechanism of aggregate formation by assessing the interaction of bacterial adhesins with synovial fluid polymer, fibrinogen. Following incubation in each simulated post-operative joint condition, the aggregates were imaged using confocal microscopy. Our analysis revealed the formation of two distinct aggregate phenotypes dependent on whether the incubation was conducted under static or dynamic conditions. Using a surface adhesin mutant, we have narrowed down the genetic determinants for synovial fluid aggregate formation and identified essential host polymers required. We report here that synovial fluid-induced aggregation is influenced by various changes specific to the post-surgical joint environment. While we now have evidence that select synovial fluid polymers facilitate bridging aggregation through essential bacterial adhesins, we suspect that their utility is limited by the increasing viscosity under static conditions. Furthermore, dynamic fluid movement recovers the ability of the bacteria with present surface proteins to aggregate under high viscosity conditions, yielding large, globular aggregates.

**Importance:** Infection is a major complication of knee and hip joint replacement surgery which is used to treat arthritis or joint damage. We have shown that *Staphylococcus aureus*, a common bacterial pathogen, aggregates upon contact with synovial fluid. Within seconds, the bacterial cells will interact with synovial fluid polymers in the joint fluid through their cell wall adhesins. The rapid formation of these aggregates likely aids in early bacterial survival in the joint-potentially contributing to the likelihood of developing an infection. By strengthening our basic understanding of the mechanics of synovial fluid aggregate formation under clinically relevant conditions, we hope to expand the knowledge of how to prevent or disrupt aggregation and reduce and more successfully treat these joint infections.

## Introduction

In recent years, synovial fluid-induced aggregation of *Staphylococcus aureus* has been heavily investigated as a bacterial survival mechanism in the development of periprosthetic joint infections (PJI). Numerous *in vitro* studies have provided novel insights into both the composition and pathogenic attributes of staphylococcal aggregates(1),(2),(3),(4),(5). It is now evident that aggregation occurs rapidly upon contact with synovial fluid and confers considerable recalcitrance to antibiotic administration (1),(3),(6)(7). To translate these hallmark findings to the clinic, it is critical that we first understand the primary mechanisms which mediate synovial fluid-induced aggregation as well as how they are influenced by the post-surgical joint environment.

Bacterial aggregates have been observed existing both within the synovial fluid of chronically infected patients as well as recapitulated in the laboratory, often with the use of bovine, porcine, or equine synovial fluid (3)(8). In previous work, we demonstrated that synovial fluid-induced aggregation can be stimulated by individual components of synovial fluid, specifically fibrinogen and fibronectin (1). More recently, a synthetic synovial fluid composed of fibrinogen, hyaluronic acid, and albumin was shown to facilitate the formation of staphylococcal aggregates *in vitro* (4). These studies indicate that bacterial binding to host polymers within the synovial fluid plays a significant role in the development of synovial fluid-induced aggregates. However, it is still unclear how localized changes within the synovial joint in the context of surgery influence this process.

The synovial joint environment during and following surgery is highly dynamic, with several distinguishing characteristics from a native joint. Following surgical site closure, invading bacteria will encounter synovial fluid that is in compositional flux. The infiltration of blood into the joint dilutes the overall concentration of synovial fluid polymers which are suspected to facilitate aggregate formation (9). Furthermore, localized inflammation without indication of infection triggers a rapid influx of immune cells, leading to both protein degradation and hyaluronic acid cleavage (10),(11),(12),(13),. Hyaluronic acid, a glycosaminoglycan produced by resident synoviocytes, provides synovial fluid with its viscous property-essential for proper joint function (14),(15),(16). The breakdown of hyaluronic acid in the inflamed joint decreases the overall viscosity of the synovial fluid (17).

After a surgical procedure, the joint cavity will be predominantly filled with blood. Over time, host cells lining the synovial membrane secrete synovial fluid back into the cavity, subsequently displacing the blood and changing the rheological properties of the fluid (9),(18). Together, the post-surgical joint becomes a complex system of shifting factors that potentially influence aggregation. We generated and tested several hypotheses based on current findings in the literature. Because synovial fluid-induced aggregation is dependent on binding to protein polymers within the fluid, we hypothesized that a compositional flux (e.g., a refilling post-surgical joint) would influence aggregate formation. We suspected that the addition of synovial fluid would enhance aggregation in a dose-dependent manner, with more circulating polymers providing a greater potential for bacterial bridging (19).

Following incubation in different concentrations of bovine synovial fluid (BSF), the average aggregate size and branch lengths were quantified using confocal microscopy followed by image analysis software. As the overall viscosity changes over time due to both surgery-induced inflammation and synovial fluid infiltration, we next evaluated the influence of viscosity on aggregation, independent of increasing synovial fluid polymers. Finally, the post-operative synovial joint is seldom a static environment, with joint motion encouraged immediately following surgery. Inducing joint flexion will stimulate the movement of synovial fluid and blood in the joint, creating a dynamic environment (20). To investigate the effect of fluid dynamics and shear stress on the formation of synovial fluid aggregates, the bacteria were imaged macroscopically after incubation on either an orbital shaker or a rocker.

During our study, we noticed distinct phenotypic differences between aggregates formed under static and dynamic conditions. As both conditions will likely exist at some point following surgical site closure, we wanted to understand the mechanics of aggregate formation under both conditions. With the use of *S. aureus* quadruple surface adhesin mutant, AH4413 *(ΔclfA ΔfnbAB clfB: Tn*), we probed the necessity of fibrinogen and fibronectin-binding proteins under static incubation as well as on a shaker. Taken together, this work demonstrates that the unique characteristics of the post-surgical joint environment-including changes in fluid dynamics, synovial fluid concentration, and viscosity influence early aggregation-potentially contributing to the efficacy of synovial fluid-induced aggregation as a survival mechanism. Furthermore, we report evidence of a polymer bridging mechanism for aggregate formation which requires the presence of specific bacterial surface adhesins.

## Results

### Staphylococcal aggregates rapidly grow and adopt a branching phenotype over time following synovial fluid contact in a static system

We previously reported that *S. aureus* will rapidly aggregate upon contact with bovine synovial fluid (BSF) (1). With this preliminary knowledge, we first sought to validate our image-analysis-based method for quantifying aggregate size and branching. Throughout a 1-hour incubation in BSF, 5 representative images of the aggregates were captured at various time points. Microscopy revealed a relative increase in aggregate size over time (**Fig. 1a**) with the average aggregate increasing by approximately 2µm^2^ over one hour (**Fig. 1b**). Time-lapse videos were collected in order to visualize the early kinetics of aggregate formation upon contact with PBS (**Supplemental Movie 1**), 10% BSF in PBS (**Supplemental Movie 2**), or 10% human serum in PBS (**Supplemental Movie 3**). Aggregate branching was also assessed to quantify aggregate morphology, which can differentiate mechanisms of polymer-mediated aggregation. Using image analysis software, we quantified the 50 longest aggregate branches at parallel time points throughout the one-hour incubation (**Fig. 2a**). Similar to aggregate size, branch length also correlated positively to BSF exposure time (**Fig. 2b**).

**Figure 1.**
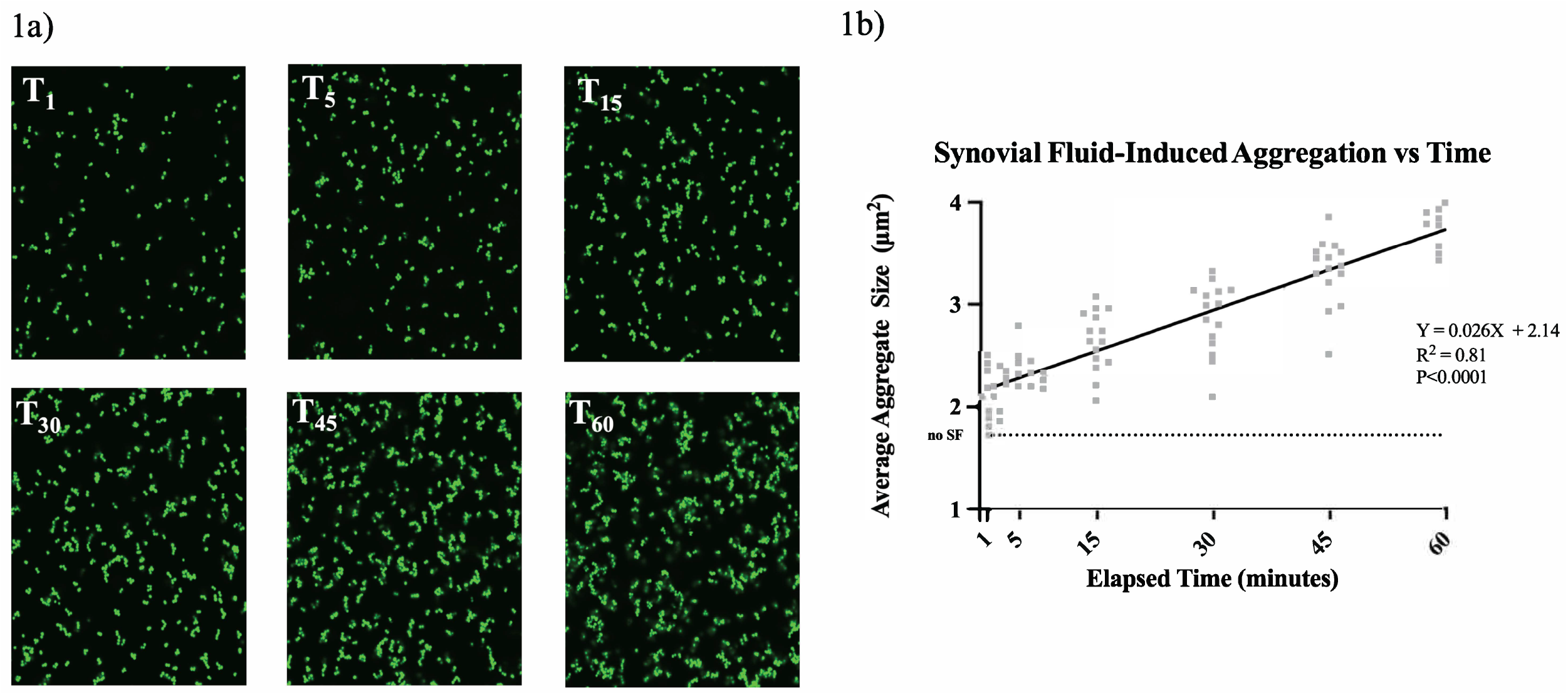
Aggregation of *Staphylococcus aureus* following contact with bovine synovial fluid (BSF). Confocal imaging of GFP-expressing *S. aureus* began immediately upon contact with BSF and continued for 1-hour with 5 representative images captured at 0, 1-, 5-, 15-, 30-, 45-, and 60-minute time points (**1a**). FIJI image analysis software was used to quantify the aggregate size at each point (**1b**). Additionally, continuous time-lapse videos were conducted to capture the initial kinetics of aggregate formation following bacterial contact with either PBS (**1c**), 10% BSF in PBS (**1d**), or 10% human serum in PBS (**1e**). Images were captured every minute for 1-hour at 60x magnification with an additional 2x zoom. To determine whether the relationship between time and aggregate size is statistically significant, a linear regression analysis was performed, indicated by the black regression line. Dashed line indicates the average size of untreated bacteria after 1-hour. Data points indicate 3 biological replicates.

**Figure 2.**
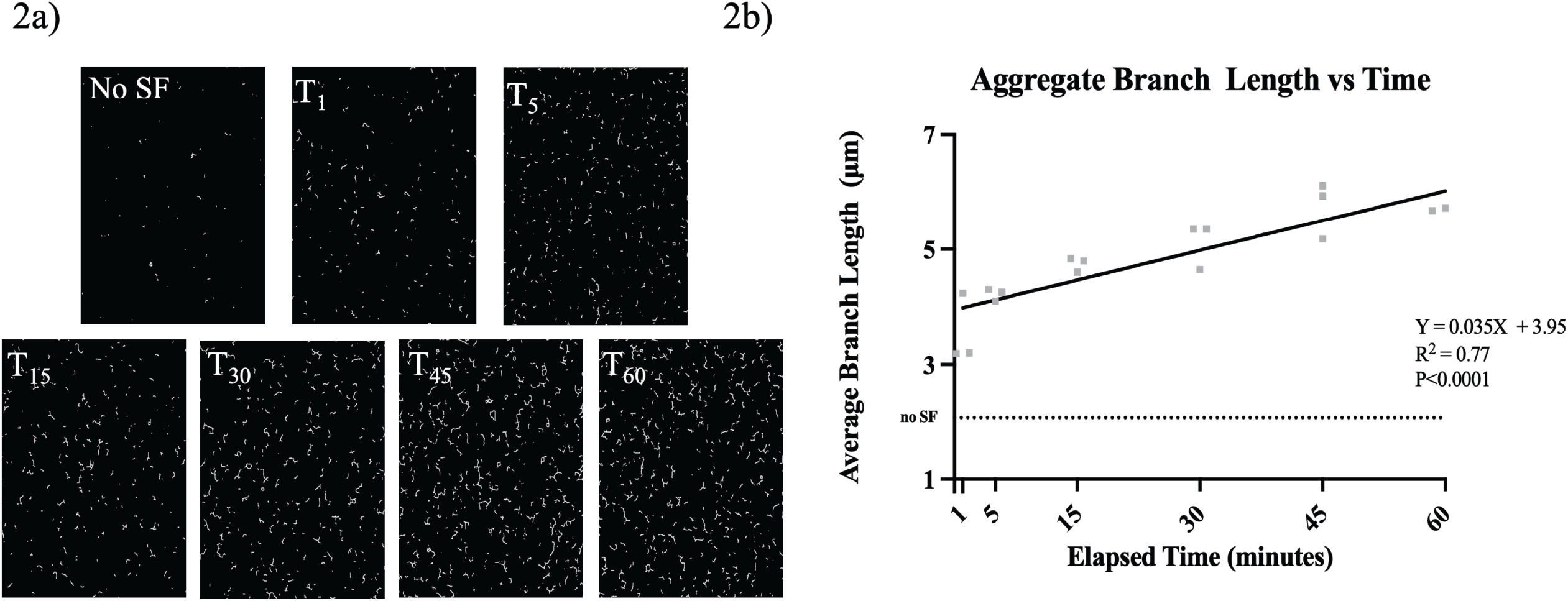
Branching analysis of *Staphylococcus* aureus following contact with bovine synovial fluid (BSF). Confocal imaging was conducted immediately upon contact with BSF and continued for 1-hour with 5 representative images collected at 0, 1-, 5-, 15-, 30-, 45-, and 60-minute time points. FIJI image analysis software was used to skeletonize the aggregates for branching quantification (**2a**). The average branch length of the 50 longest branches was calculated for each time point (**2b**). To determine whether the relationship between time and branch length is statistically significant, a linear regression analysis was performed, indicated by the black regression line. Dashed line indicates the average branch length of untreated bacteria after 1-hour. Data points indicate the average of 3 biological replicates.

### Synovial fluid induces an aggregate size threshold at high concentrations

Using our image analysis methodology, we next quantified aggregate size and branching as a function of various surgical joint-specific exposures. Previous studies show that binding to protein polymers within the synovial fluid is essential for synovial fluid-induced aggregation of *S. aureus* to occur (20),(4). We hypothesized that increasing the synovial fluid concentration and thus increasing the concentration of available bridging polymers, would facilitate the formation of larger aggregates with longer branches. As anticipated, we observed a relative increase in the average aggregate size after raising the synovial fluid concentration from 10% BSF to 20% BSF (**Fig. 3a and Fig. 3b**). Interestingly, increasing the concentration to 50% BSF resulted in both a reduction in the average aggregate size and branch length (**Fig. 3c and Fig. 3d**) These findings suggest that the increase in viscosity associated with the higher concentrations may inhibit synovial fluid polymer binding as well as inter-bacterial interactions.

**Figure 3.**
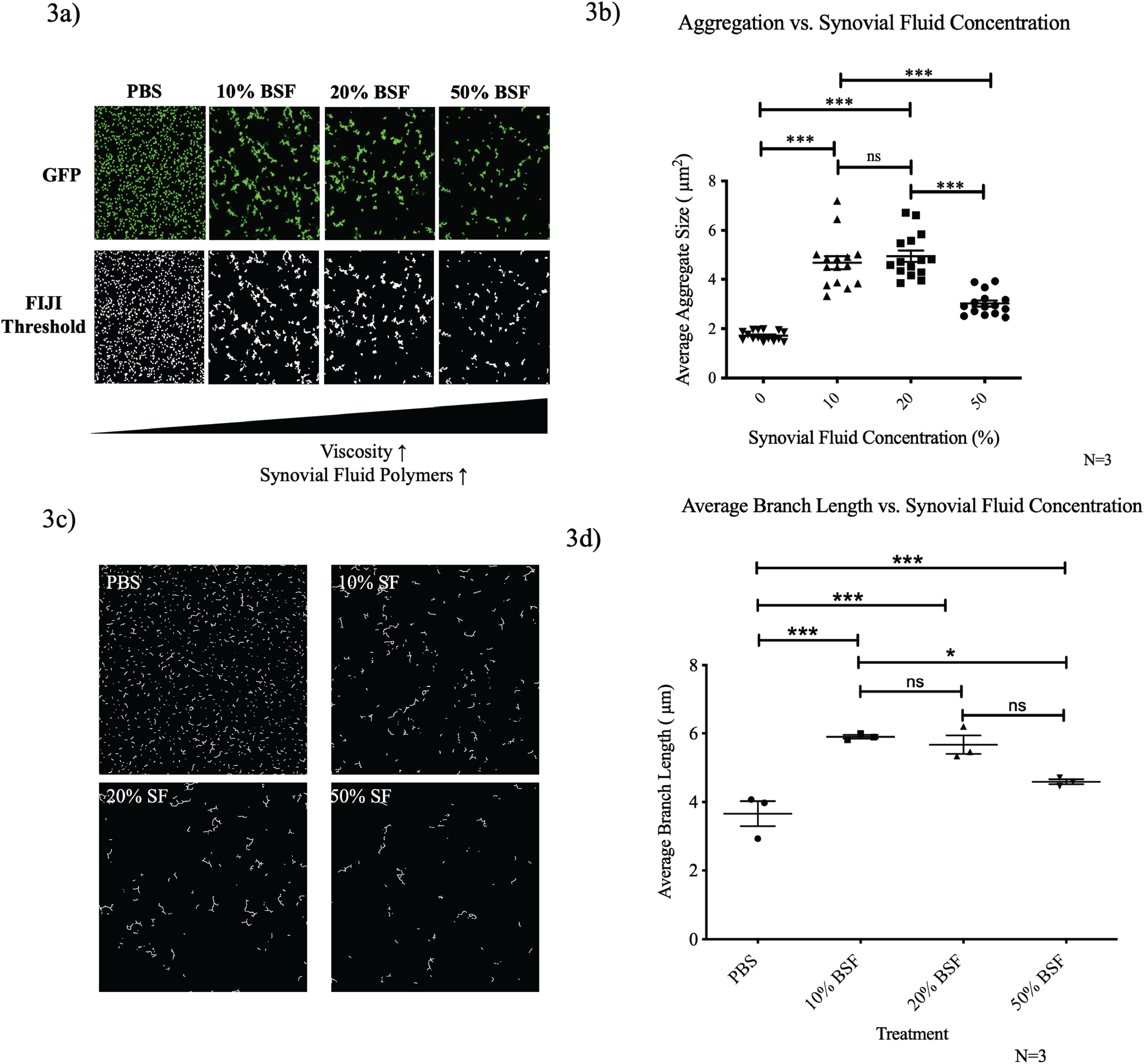
Increasing the concentration of bovine synovial fluid (BSF) inhibits aggregation under static conditions. Synovial fluid-induced aggregates were imaged using confocal microscopy following 1-hour of incubation in either PBS, 10% BSF, 20% BSF, or 50% BSF (**3a**). FIJI image analysis software was used to quantify the average particle size (**3b**). Images were skeletonized (**3c**) and the average branch length of the 50 longest branches was calculated (**3d**). Data points represent 3 biological replicates with 5 representative images collected for each replicate. Error bars indicate mean +/- SEM. Statistical significance was determined by one-way ANOVA by a Bonferroni’s Multiple Comparison Test to compare means between treatments. ns p>0.05; *** p ≤ 0.001.

### Increasing solution viscosity with hyaluronic acid inhibits staphylococcal aggregation independent of synovial fluid protein polymers

To investigate the effect of an increasingly viscous joint space on synovial fluid-induced aggregate formation, various concentrations of purified, high molecular weight hyaluronic acid were supplemented into a baseline concentration of 10% BSF. Hyaluronic acid was chosen because it is the primary contributor to the viscous property of synovial fluid and does not stimulate staphylococcal aggregation independent of the other synovial fluid components at physiological concentrations (1). Prior to imaging studies, we first measured the viscosity of the different synovial fluid concentrations and hyaluronic acid-supplemented synovial fluid using a rheometer. As we increased the concentration of synovial fluid in the solution, the viscosity increased in a dose-dependent manner (**Fig. 4a**). Similarly, supplementation with purified hyaluronic acid elevated the overall viscosity (**Fig. 4b**). Increasing the concentration of hyaluronic acid on top of the baseline 10% BSF decreased both the average size (**Fig. 4c and Fig. 4d**) of the aggregates as well as the length of the branches (**Fig. 4e** and **Fig. 4f**), indicating that viscosity plays a significant role in the extent of synovial fluid-induced aggregation.

**Figure 4.**
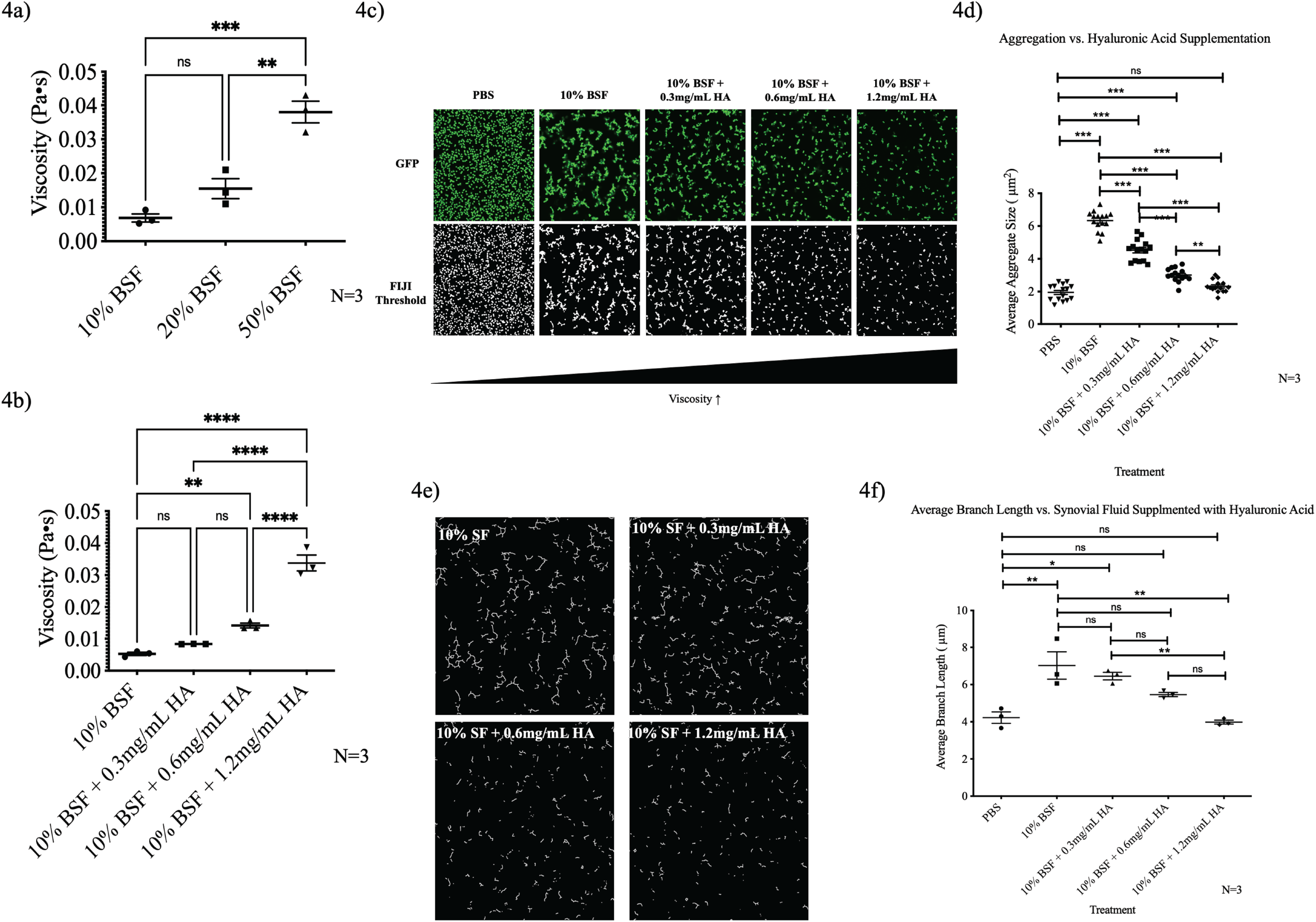
Increasing the solution viscosity inhibits synovial fluid-induced aggregation independent of increasing fibrinogen and fibronectin bridging polymers. A rheometer was used to measure the viscosity of increasing concentrations of bovine synovial fluid (BSF) (**4a**) as well as 10% BSF supplemented with increasing concentrations of hyaluronic acid (**4b**). Synovial fluid-induced aggregates were imaged using confocal microscopy following 1-hour of incubation in either PBS, 10% BSF, or 10% BSF supplemented with various concentrations of hyaluronic acid (**4c**). FIJI image analysis software was used to quantify the average particle size (**4d**). Images were skeletonized (**4e**) and the average branch length of the 50 longest branches was calculated (**4f**). Data points represent 3 biological replicates with 5 representative images collected for each replicate. Error bars indicate mean +/- SEM. Statistical significance was determined by one-way ANOVA by a Bonferroni’s Multiple Comparison Test to compare means between treatments. ns p>0.05; ** p ≤ 0.01; *** p ≤ 0.001.

### Dynamic incubation of *S. aureus* in synovial fluid recovers aggregation at high synovial fluid concentrations

In addition to changes in synovial fluid viscosity and polymer composition within the post-surgical joint cavity, there will also be the introduction of fluid flow (20). Synovial fluid and blood flow within the enclosed space will be stimulated by normal joint flexion as well as prescribed therapeutic flexion (21). To probe the influence of fluid dynamics on synovial fluid-induced aggregation of *S. aureus*, we incubated the bacteria statically, on an orbital shaker, or a rocker, for 1-hour in various concentrations of BSF before macroscopic imaging. Under static conditions, macroscopic aggregates were not visible in the Petri dishes (**Fig. 5a**). However, under dynamic conditions free-floating, globular aggregates were observed, even at high concentrations of synovial fluid (**Fig. 5b**). Macroscopic imaging was also conducted following incubation on a rocker which we expect more closely resembles joint flexion-induced fluid flow. Similar to the orbital shaker, the rocker yielded large aggregates; however, they were less globular in morphology (**Fig. 5c**). Finally, macroscopic aggregates were imaged after incubating in increasing concentrations of hyaluronic acid (**Fig. 5d**). These findings indicate that fluid dynamics can overcome the negative effect of viscosity on aggregation as previously observed.

**Figure 5.**
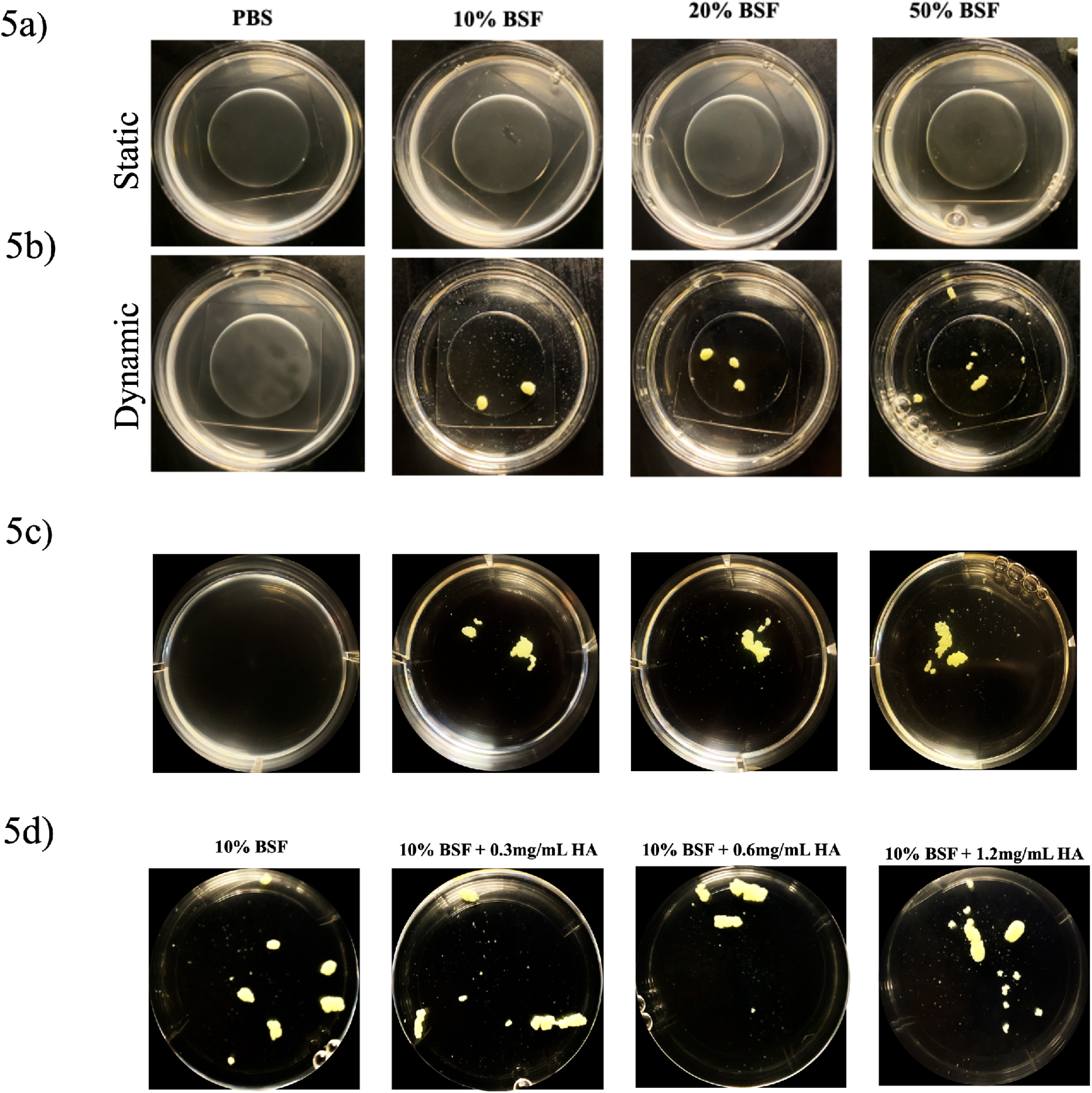
Dynamically incubating *Staphylococcus aureus* in bovine synovial fluid (BSF) recovers aggregate formation at high concentrations of synovial fluid and high viscosity solutions. *S. aureus* following was imaged following 1-hour of incubation statically on a benchtop, (**5a**) dynamically on an orbital shaker, (**5b**) or a rocker (**5c**) with increasing concentrations of BSF. Additionally, macroscopic imaging was conducted following dynamic incubation on an orbital shaker with increasing concentrations of hyaluronic acid (**5d**). Following a 1-hour incubation, Petri dishes were imaged macroscopically with a camera.

### Fibronectin-binding proteins and clumping factors mediate rapid aggregation through a polymer bridging mechanism

As the presence of fibrinogen and fibronectin is reportedly essential for bacterial aggregation in synovial fluid, we next sought to validate the importance of select microbial surface components recognizing adhesive matrix molecules (MSCRAMMs) using a *S. aureus* quadruple adhesin mutant strain *(ΔclfA ΔfnbAB clfB::Tn*). Following 1-hour of static incubation in either PBS or 10% BSF in PBS, both the wild type *S. aureus* and adhesin mutant were imaged using confocal microscopy (**Fig. 6a**). While both strains were capable of forming phenotypically comparable branched aggregates, image analysis quantification revealed that the adhesin mutant formed significantly smaller aggregates compared to those of the wild type (**Fig. 6b**) with a higher degree of aggregate circularity (**Fig. 6c**). When incubated under dynamic conditions on an orbital shaker, the wild-type strain conglomerated into a macroscopic, free-floating aggregate, while the mutant strain remained dispersed throughout the Petri dish (**Fig. 6d**).

**Figure 6.**
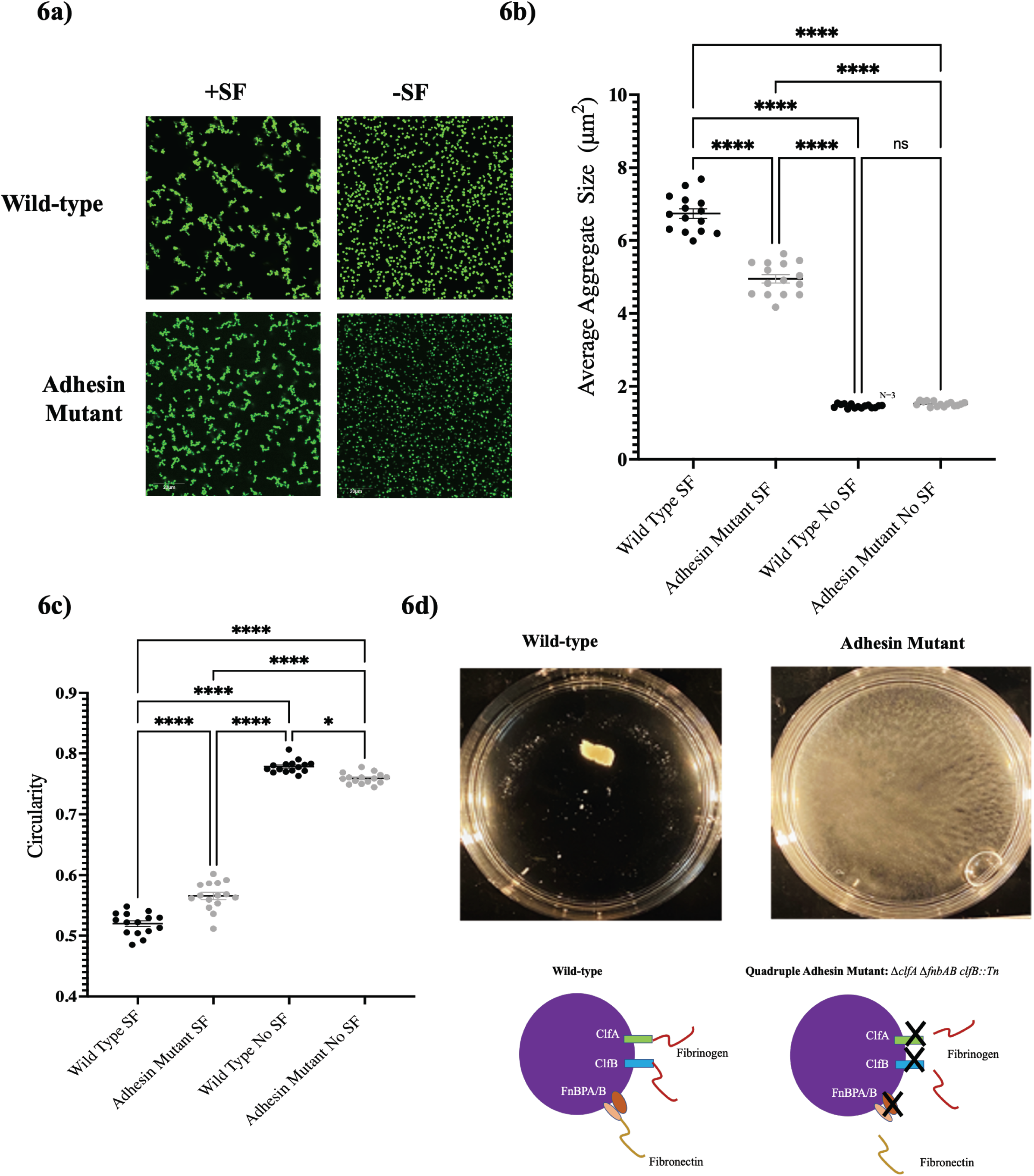
Key surface adhesins are essential for macroscopic aggregation under dynamic incubation but not required for the static formation of microscopic branching aggregate phenotype. Wild type *Staphylococcus aureus,* AH1263, and quadruple adhesin mutant, AH4413 (*ΔclfA ΔfnbAB clfB::Tn*) were incubated statically in 10% bovine synovial fluid or PBS before imaging with confocal microscopy (**6a**). The average aggregate size (**6b**) and circularity (**6c**) were determined using FIJI image analysis software after 1-hour of incubation. Following 1-hour of dynamic incubation on an orbital shaker, the aggregates were imaged macroscopically (**6d**). Error bars indicate mean +/- SEM. Data points represent 3 biological replicates with 5 representative images collected for each replicate. Error bars indicate mean +/- SEM. Statistical significance was determined by one-way ANOVA by a Bonferroni’s Multiple Comparison Test to compare means between treatments. ns p > 0.05; ** p ≤ 0.01; *** p ≤ 0.001, ****p ≤ 0.0001

We hypothesized that this defect was due to a lost interaction between one or more bacterial surface adhesins and the synovial fluid polymer. We repeated the experiment using purified Alexa-Fluor conjugated fibrinogen. Fibrinogen alone was selected as it has previously been documented to play an important role in staphylococcal aggregation (1),(4). Similar to the 10% BSF, the wild-type strain formed a single, macroscopic, free-floating aggregate following 1-hour of dynamic incubation on a shaker. Upon microscopic analysis, we visually observed strong fibrinogen co-localization. In contrast, the adhesin mutant was unable to form a free-floating aggregate under shear and displayed little interaction with the fibrinogen (**Fig. 7a**). Interestingly, under static conditions, the adhesin mutant was unable to form the branched aggregates which were still observed following static incubation in the 10% BSF (**Fig. 7b**).

**Figure 7.**
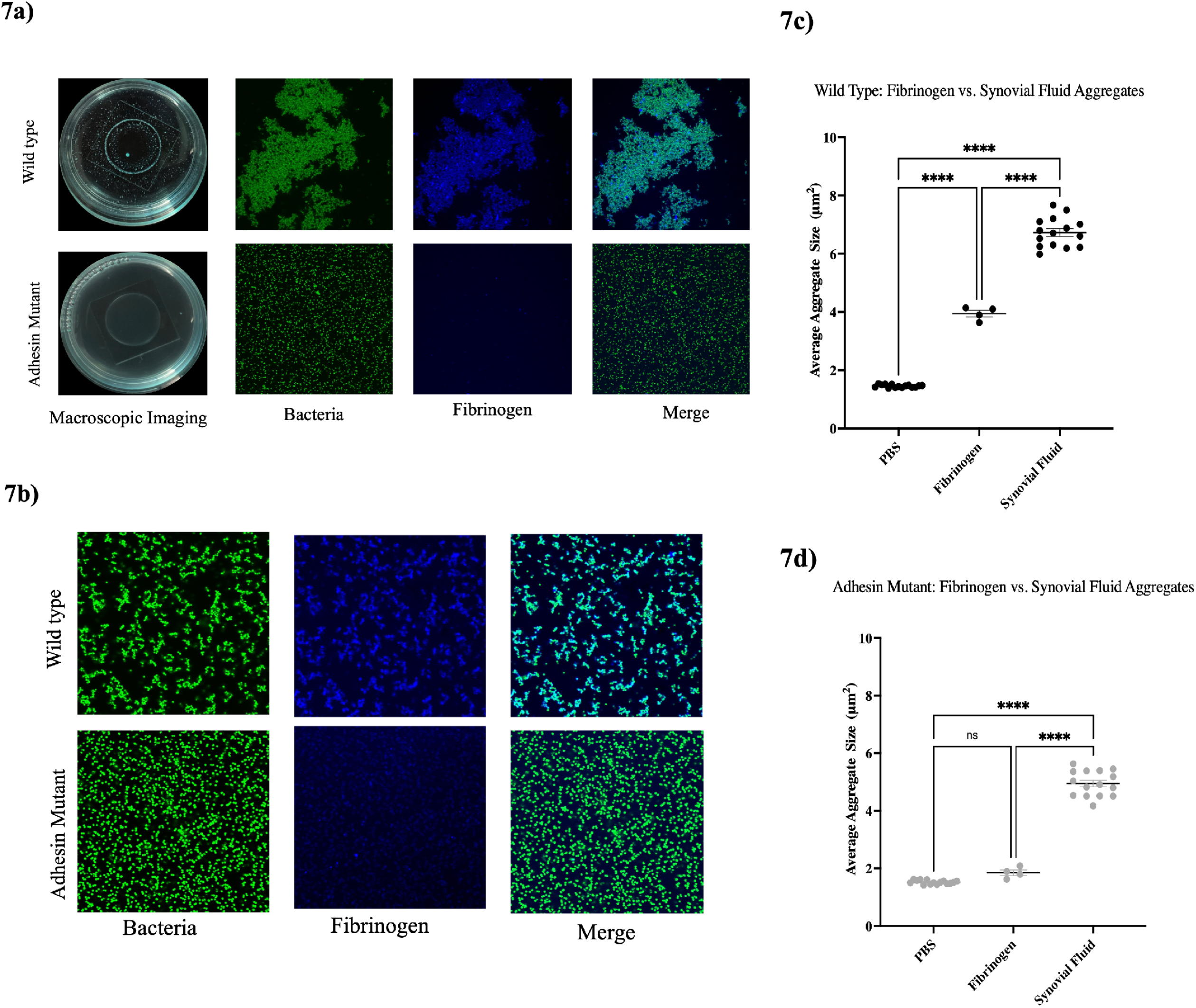
Polymer bridging aggregation is facilitated by direct interaction between bacterial surface adhesins and fibrinogen. Macroscopic and confocal microscopy images were captured of both wild-type *Staphylococcus aureus* and the quadruple adhesin mutant following 1-hour of either dynamic (**7a**) or static (**7b**) incubation in Alexa Fluor-conjugated fibrinogen. Following static incubation, the average aggregate size was calculated for the wild type (**7c**) and mutant (**7d**). Error bars indicate mean +/- SEM. Data points represent 3 biological replicates with 5 representative images collected for each replicate. Error bars indicate mean +/- SEM. Statistical significance was determined by one-way ANOVA by a Bonferroni’s Multiple Comparison Test to compare means between treatments. ns p > 0.05; ** p ≤ 0.01; *** p ≤ 0.001, ****p ≤ 0.0001

Quantification through image analysis revealed that fibrinogen alone produced aggregates that were 2-fold larger than the untreated wild-type bacteria and 2-fold smaller than the 10% BSF aggregates (**Fig. 7c**). There was no significant difference between the untreated adhesin mutant and that treated with the fibrinogen (**Fig. 7d**). Experiments with the adhesin mutant reveal that mutations in select MSCRAMMs abolish the aggregate phenotype under dynamic conditions but not static ones. Statically stimulated aggregation of the mutant is ultimately lost when incubated with a single synovial fluid polymer, fibrinogen. As the wild type is still able to aggregate in fibrinogen alone, this suggests a direct role for fibrinogen in synovial-fluid induced aggregation as well as an interaction, at the genetic level, between fibrinogen and one or more MSCRAMMs. Furthermore, the ability of the adhesin mutant to still aggregate statically in the 10% BSF but not the purified fibrinogen indicates other polymers within the synovial fluid may be contributing independently to the branched phenotype.

To solidify our claim that synovial fluid-induced aggregation is mediated by a rapid bridging interaction between bacterial surface proteins and host polymers- as opposed to an active process- we conducted aggregation assays with antibiotic-treated bacteria. Following treatment with gentamicin, non-viable *S. aureus* was incubated in 10% BSF and assessed for aggregation. Confocal imaging revealed that throughout a 1-hour incubation, the dead bacteria were still able to aggregate in the 10% BSF (**Supplemental Fig. 3a**). These data indicate that the presence of the surface adhesins is sufficient to facilitate aggregate formation and further explain the rapidity with which the bacteria come together upon synovial fluid exposure.

### Dynamic bacterial aggregation becomes less efficient as synovial displaces blood

As mentioned previously, following surgical site closure, blood will predominantly occupy the joint cavity. To mimic the influence of the transition from a blood-dominated cavity to a synovial fluid-dominated one, we conducted macroscopic imaging after 1-hour of dynamic incubation in synovial fluid supplemented with human blood (**Fig. 8a**). Comparable aggregation was observed between the 100% blood condition and the combination of 90% blood with 10% BSF condition. Interestingly, at higher concentrations of BSF (<75%), macroscopic aggregation appeared dwarfed, with smaller bacterial aggregates and clusters forming. Confocal microscopy of the 10% BSF in blood was conducted to confirm the observed aggregates were GFP-tagged staphylococcal aggregates (**Fig. 8b**).

**Figure 8.**
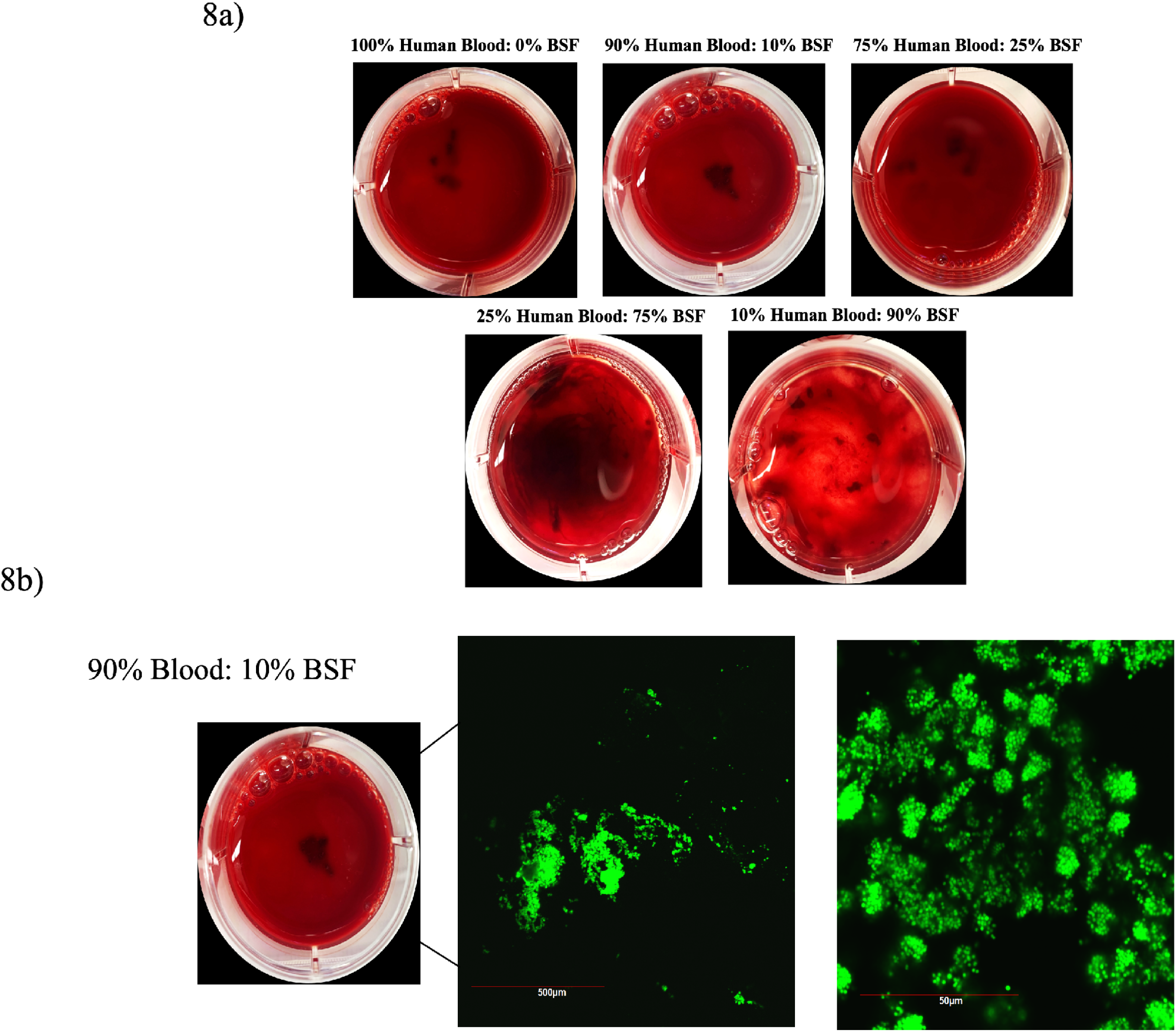
Dynamic incubation of *Staphylococcus aureus* in human blood supplemented with bovine synovial fluid (BSF). GFP-expressing *S. aureus* was incubated for 1-hour in human blood supplemented with increasing concentrations of BSF. Macroscopic imaging of the Petri plates was conducted of the free-floating aggregates (**8a**) followed by confocal microscopy to validate the presence of bacteria in the aggregates (**8b**).

## Discussion

Synovial fluid-induced aggregation of *S. aureus* is under investigation as a potential pathogenic survival mechanism in the occurrence of periprosthetic joint infections. While a considerable amount of work has been conducted characterizing this process, until now, the influence of surgical site-specific conditions on aggregate formation has been neglected. Following prosthetic joint surgery, numerous changes associated with inflammation and joint flexion will occur. These changes are highly distinct from a native joint environment, and as such, should be taken into consideration when studying the establishment and progression of bacterial infections. Furthermore, analysis of synovial fluid-induced aggregates has been confined to long incubations (<90 minutes), leaving the initial kinetics of aggregation unclear.

We previously published work demonstrating that staphylococcal aggregation occurs rapidly upon contact with BSF. Using confocal microscopy, *S. aureus* was imaged immediately following exposure to either 10% BSF in PBS, 10% human serum in PBS, or PBS alone. Our videos reveal that not only does aggregation occur immediately upon contact with the BSF, but the resulting aggregates form distinctive branched structures as they develop. In contrast to tightly packed, globular bacterial clusters which form from polymer depletion, the branching structures we observed appear visually similar to those generated through diffusion- and reaction-limited cluster aggregation (DLCA and RLCA) (22). This model was developed from colloidal physics to explain the shape and growth of different structures formed from colloidal particles within a liquid. In this case, particles will diffuse through a surrounding medium and stick together irreversibly whenever they collide (DLCA), or with a certain probability upon each collision (RLCA) (22).

Two dominating mechanisms of bacterial aggregation when either host-produced or bacterial polymers are present, have recently been proposed (19). Depletion aggregation is an entropy-driven process, in which high concentrations of non-adsorbing polymers constrain the bacteria together. This mechanism predominantly yields tightly packed bacterial aggregate arrangements due to spontaneous motion of the surrounding polymers-pushing the bacteria together and maximizing the space in which the polymers have to move (19). The second primary mechanism of polymer-induced aggregation is bridging aggregation, where the host or bacterial polymers adsorb on the bacterial cells (19).

While synovial fluid is a complex host fluid composed of numerous polymers, we speculated that synovial fluid-induced aggregation of *S. aureus* is dominated by the bridging aggregation mechanism, with fibrinogen or fibronectin serving as bridging polymers (19). The highly branched morphology of the synovial fluid aggregates corroborates this theory, as bridging aggregation yields random bacterial arrangements due to the attachment of polymers interacting with various surface adhesins dispersed across the cell surface (19). In contrast, the human serum aggregates also grew over time but revealed less aggregation with fewer intricate branches. These results were expected as serum lacks fibrinogen and fibronectin polymers required for extensive bridging aggregation, but still contains albumin at high concentrations, likely facilitating the observed clumping.

Based on our observations from the time lapse microscopy of the initial kinetics, we hypothesized that aggregate size and branching would increase with synovial fluid concentration. To assess the influence of various synovial fluid concentrations on aggregate formation, we incubated *S. aureus* for 1-hour in PBS, 10% BSF in PBS, 20% BSF in PBS, and 50% BSF in PBS before confocal imaging. To our surprise, both aggregate size and branch length increased in a dose-dependent manner with the synovial fluid concentration up until the 50% BSF treatment. Although the amount of available bridging polymers elevates with the synovial fluid concentration, the overall viscosity of the solution also increases. These results suggest that a threshold exists at which the synovial fluid is too viscous for inter-bacterial interaction. We suspect that once this threshold is reached, even an abundance of fibrinogen and fibronectin cannot recover the larger aggregates under static conditions. This data indicates that the early time points following surgical site closure, when the blood to synovial fluid ratio is higher, may facilitate the most optimal conditions for staphylococcal aggregate formation.

To probe our theory that a viscosity-induced threshold for aggregation exists at higher concentrations of synovial fluid, we quantified both aggregate size and branching following incubation in 10% BSF supplemented with increasing concentrations of high molecular weight hyaluronic acid sodium salt. The addition of hyaluronic acid increased the viscosity of the solution dose-dependently and inhibited aggregation of *S. aureus* independent of increasing concentrations of synovial fluid protein components. We acknowledge that the use of hyaluronic acid as an agent for viscosity manipulation introduces a confounding variable into our system. However, previous studies in our laboratory show that hyaluronic acid alone does not stimulate staphylococcal aggregation at the applied concentrations (1).

While a static synovial joint environment may closely mimic the immediate conditions following surgical site closure, most surgeons advise patients to introduce movement as part of a standard post-operative protocol (21). Not only are patients fully weight-bearing after surgery, but they are encouraged to partake in physical therapy to maintain a full range of motion, including maximal knee flexion. Induced synovial fluid movement introduced post-operatively likely creates a highly interactive environment, enhancing inter-bacterial interaction as well as increased contact with circulating synovial fluid polymers. With the use of an orbital shaker, we incubated *S. aureus* dynamically for 1-hour in either PBS, 10% BSF in PBS, 20% BSF in PBS, or 50% BSF in PBS. Macroscopic imaging revealed the presence of large, free-floating aggregates, even at the higher concentrations of BSF. These findings suggest that dynamic synovial fluid movement recovers the ability of the bacteria to aggregate under conditions of high viscosity, as it facilitates a greater degree of bacterial interaction with each other and the bridging polymers. We also conducted dynamic aggregation experiments on a rocker, as the motion will more closely resemble the back-and-forth movement stimulated by joint flexion (20). While the rocker-induced aggregates were less globular in morphology, the apparatus was still capable of stimulating synovial fluid aggregates at each concentration of synovial fluid. It is possible that the introduction of fluid flow may lower the bacterial concentrations required to stimulate aggregate formation as it will enhance the probability of collisions.

As previously described, the synovial cavity following joint surgery will primarily be filled with blood. Human blood is approximately twice as viscous as PBS and contains excess polymers within the serum and plasma components which may contribute to early aggregate formation (23),(24). To simulate the refilling of synovial fluid into the joint space whilst displacing the blood, we incubated *S. aureus* for 1-hour on a shaker with various concentrations of synovial fluid and heparinized human blood. Although heparin is seldom used during surgery, patients are often placed on anticoagulants either the day of surgery or post-operatively. As expected, we noticed a degree of macroscopic aggregate formation in the 100% blood. Supplementation of either 10% BSF or 25% BSF into the blood produced comparable-sized macroscopic aggregates. However, higher concentrations of synovial fluid with low amounts of blood diminished the large, globular aggregate phenotype. We suspect that in combination with the blood, higher concentrations of synovial fluid create a system too viscous to facilitate aggregate formation, even under dynamic conditions. This data corroborates our hypothesis that early time points consisting of lower synovial fluid concentrations may be optimal for aggregate formation. Interestingly, upon microscopic observation, our blood-10% synovial fluid aggregates displayed phenotypic similarity to synovial fluid aggregates observed in the synovial fluid of patients with infected joints (8). Our findings indicate that not only is the presence of synovial fluid polymers important for aggregate formation but that their utility by *S. aureus* is highly influenced by the state of the joint environment.

Following surgical site closure, there will likely be periods of both static as well as dynamic fluid conditions which stimulate the formation of distinct aggregate phenotypes. As such, we sought to understand the mechanics of synovial fluid-induced aggregation under both conditions. Because the presence of fibrinogen and fibronectin has been widely reported to be essential for aggregate formation in synovial fluid, we first tested whether direct bacterial interaction with these polymers was required to form both aggregate phenotypes. With the use of a fibronectin-binding protein and clumping factor quadruple mutant strain, we examined the necessity for key binding proteins in synovial fluid aggregate formation under both dynamic and static conditions.

While the mutant strain was capable of aggregating under static conditions to a degree, under dynamic conditions, the macroscopic, free-floating aggregate phenotype was inhibited. These findings support our hypothesis that aggregation in synovial fluid is potentially a multi-step process, with requirements changing under different joint conditions. While it is likely that the adhesin mutant strain is still capable of weakly binding these polymers using other binding factors, our observations indicate that they are insufficient for forming stable macroscopic aggregates under shear (25).

To directly assess the requirement of fibrinogen binding by the bacterial surface adhesins, microscopy and macroscopic imaging were conducted following both dynamic and static incubation in 0.2 mg/mL of Alexa Fluor 488 conjugated human fibrinogen. In contrast with incubation in the 10% BSF, the adhesin mutant was unable to form either a macroscopic aggregate under dynamic conditions or branched aggregates under static conditions, while fibrinogen was sufficient under both conditions to stimulate both phenotypes with the wild type strain.

Interestingly, gentamicin-killed bacteria also aggregated upon contact with 10% BSF, indicating that active bacterial processes are not required for synovial fluid-induced aggregation to occur and that the presence of the surface adhesins is sufficient. Unexpectedly, gentamicin-killed bacterial aggregates were on average 20% larger than the live bacteria in the untreated control. This may be due to changes in bacterial expression in response to antibiotic stress. It has been reported that within 30 minutes of gentamicin exposure (100x the MIC), there is a significant elevation in the expression of SigB- a stress-responsive alternative sigma factor (26),(27). SigB activity positively influences the expression of clumping factors and fibronectin-binding proteins, which could explain the increase in aggregation observed with the gentamicin-treated bacteria (28),(29),(30). Taken together, these findings confirm our speculation that bridging aggregation does occur through the rapid interaction of bacterial adhesins with fibrinogen but suggest that it alone is not the only contributor to the aggregate phenotype.

A possible explanation for our observation of static aggregation in 10% BSF but not fibrinogen alone, is the capacity for the mutant to interact with or be influenced by other synovial fluid polymers, such as hyaluronic acid and albumin (31),(32),(33),(34). Preliminary data shows that bovine serum albumin (BSA) stimulates clumping in both the mutant as well as the wild-type strain. Furthermore, high molecular weight hyaluronic acid has been documented to facilitate depletion aggregation in *Pseudomonas aeruginosa*-yielding compact, aligned bacterial aggregates (19). The domination of the depletion mechanism in the absence of functional polymer bridging could explain the higher degree of circularity observed in the adhesin mutant aggregates compared to the highly branched wild type following incubation in 10% BSF. We are currently evaluating each of these possibilities by use of a synthetic synovial fluid to assess the influence of other host components on aggregate formation under each condition.

Synovial fluid-induced aggregation has recently been acknowledged as a potential contributor to the occurrence of orthopedic joint infections. Our basic understanding of aggregate formation in synovial fluid has grown exponentially over the past decade, however, future studies must be inclusive of conditions that consider the infection environment. In the context of joint surgery, the infection environment is both dynamic and inflammatory, with fluctuating concentrations of blood, synovial fluid, and the polymers which they are composed of. We report here that changes in these factors significantly impact the extent and rapidity of synovial fluid-induced aggregation of *S. aureus.* Additionally, we observe that fluid dynamics may play a critical role in aggregation, potentially facilitating aggregate formation at lower bacterial inoculums and higher synovial fluid concentrations. In addition to assessing the influence of joint conditions on aggregate formation, our work provides evidence of a polymer bridging mechanism through interaction between key staphylococcal surface adhesins and fibrinogen. As synovial fluid is a complex fluid, we hope to further tease apart the specific bacterial and host factors dictating aggregation in future studies.

## Materials and Methods

### Bacterial strains and growth conditions

Methicillin-resistant *S. aureus* LAC strain (AH1726) expressing GFP, was used for all confocal microscopy, image analysis quantification, and growth curves. For macroscopic imaging experiments, quadruple mutant AH4413 *(ΔclfA ΔfnbAB clfB: Tn*) was utilized along with the wild-type strain, AH1263 (35). Cultures were streaked on tryptic soy agar (TSA) (BD Biosciences, Heidelberg, Germany) from agar slants (Horswill Laboratory, University of Colorado School of Medicine, Aurora, Colorado). TSA plates were incubated overnight at 37°C, and a single isolated colony was used to inoculate 5mL of tryptic soy broth (TSB) (BD Biosciences) in a 15mL falcon tube. Inoculated broth cultures were grown for 18 hours at 37°C in an orbital shaker set to 200 RPM (Innova 44; New Brunswick Scientific).

### Confocal imaging of aggregates

Overnight cultures of AH1726 were grown as previously described and diluted to an OD_600_ of 0.2. 1mL of the diluted culture was pelleted at 21,000 xg for 1 minute in a centrifuge before resuspension in 1mL of PBS. The cells were then transferred into a 35 x 10 mm glass-bottom confocal dish (MatTek Corporation, Ashland, Massachusetts) containing 2mL of PBS or PBS supplemented with synovial fluid and/or hyaluronic acid. Confocal microscopy was conducted 1-hour after bacterial incubation in PBS, 10% BSF in PBS, 20% BSF in PBS, or 50% BSF in PBS at room temperature. Additionally, the effect of viscosity on synovial fluid-induced aggregation was assessed by imaging 1-hour after incubation in 10% BSF supplemented with increasing concentrations of high molecular weight hyaluronic acid sodium salt (1.5-1.8MDa) (Alfa Aesar, Haverhill, MA, USA) in PBS. pH readings were measured for each solution to ensure pH was not influencing aggregation phenotypes (**Supplemental Figure 2**). Fibrinogen aggregates were stimulated using 0.2mg/mL of purified fibrinogen from human plasma with an Alexa Fluor 488 conjugate (Invitrogen, Waltham, Massachusetts). Images were collected using an Olympus FluoView FV10i Confocal Laser Scanning Microscope under 60x magnification with an additional 2x zoom. For each condition, 3 biological replicates were imaged with 5 representative images captured for each replicate.

### Aggregate size quantification in FIJI

Following confocal imaging, green channel images displaying GFP-tagged synovial fluid-induced aggregates were imported into FIJI image analysis software (36). Replicates were converted from individual images to stacks for batch processing. The TIFF stacks were converted to 8-bit images and a threshold was applied using an auto-threshold. A size threshold detection was set to include particles greater than or equal to 9.6 pixels, or 1µm in diameter, excluding background particles that were smaller than a single bacterial cell. Particles were analyzed and the average particle size for each image was calculated. Size data was transferred to Prism GraphPad for graphing and statistical analysis. For the synovial fluid concentrations and viscosity experiments, statistical significance was determined by one-way ANOVA followed by a Bonferroni’s Multiple Comparison Test to compare means between treatments. To determine whether the relationship between time and aggregate size was statistically significant, a linear regression analysis was performed and an R^2^ value calculated.

### Aggregate skeletonization and branching analysis

As described above, the green channel images displaying GFP-tagged aggregates were imported into FIJI image analysis software for branching quantification. The TIFF stacks were first converted to 8-bit images and made binary for skeletonization (37). Binary images were skeletonized to display branching structures for each aggregate. The size threshold detection was set to include only particles greater than or equal to 9.6 pixels (1µm) in diameter, excluding background particles that were smaller than a single bacterial cell. All branch lengths were quantified for each image and exported to excel. The longest 50 branches for each image were averaged and transferred to Prism for graphing and statistical analysis. For the synovial fluid concentration data and viscosity experiments, statistical significance was determined by one-way ANOVA followed by a Bonferroni’s Multiple Comparison Test to compare means between treatments. To determine whether the relationship between time and aggregate branch length was statistically significant, a linear regression analysis was performed and an R^2^ value calculated.

### Macroscopic imaging of dynamic aggregates

Macroscopic analysis of the dynamically formed aggregates was carried out as described in previous work (38). GFP-expressing *S. aureus* strain, AH1726, was grown overnight as described above and subsequently diluted to an OD_600_ of 0.5. The bacteria were pelleted, and the supernatant aspirated before resuspension in 1mL of PBS. The 1mL of bacteria was transferred to a 35 by 10-mm petri dish and supplemented with either 1.7mL, 1.4mL, or 500µL of additional PBS. Finally, 300µL, 600µL, or 1,500µL of BSF supernatant were added to create 10% BSF, 20% BSF, or 50% BSF in PBS solutions, respectively. The Petri dishes were incubated for 1 hour on an orbital shaker set to 60 RPM at room temperature. The same conditions were used to evaluate aggregate formation on a rocker, which we suspect will more closely resemble dynamic movement in a joint cavity. Following a 1-hour incubation on either apparatus, macroscopic imaging was conducted using a dual 12-megapixel camera secured 15 cm above the specimen. In addition to PBS, macroscopic imaging was also conducted following incubation in synovial fluid supplemented human blood. Human blood was isolated from a healthy donor by intravenous puncture following IRB protocol. Pelleted bacteria were diluted to an OD_600_ of 0.5 and resuspended in blood supplemented with increasing concentrations of BSF.

### Statistical analysis

GraphPad Prism v9.20 software was used for statistical analysis of the following data. The threshold significance was set at a P-value of 0.05. All error bars indicate the standard error of the mean (SEM). Statistically significant differences were determined using the test specified in the corresponding methods sections.

## Acknowledgements

Amelia Staats was supported by The Ohio State University College of Medicine, Program for Advancing Research in Infection and Immunity Fellowship.

PS is supported by NIH R01 GM124436

ARH is supported by NIH public health service grant AI083211

## Supplemental Material Captions

**Supplemental Movie 1. Time-lapse imaging aggregate formation in PBS.** Images were captured every minute for 1-hour at 60x magnification with an additional 2x zoom.

**Supplemental Movie 2. Time-lapse imaging aggregate formation in 10% bovine synovial fluid in PBS.** Time-lapse video was conducted to capture the initial kinetics of aggregate formation following bacterial contact with BSF. Images were captured every minute for 1-hour at 60x magnification with an additional 2x zoom.

**Supplemental Movie 3. Time-lapse imaging aggregate formation in 10% human serum in PBS.** Time-lapse video was conducted to capture the initial kinetics of aggregate formation following bacterial contact with human serum. Images were captured every minute for 1-hour at 60x magnification with an additional 2x zoom.

**Supplemental Figure 1. Growth of *S. aureus* in increasing concentrations of bovine synovial fluid (BSF) in PBS.** *Staphylococcus aureus* strain, AH1726, was incubated for 3-hours in PBS, 10% BSF in PBS, 20% BSF in PBS, or 50% BSF in PBS. Incubation was conducted shaking at 37℃ with sampling for CFU-plating every hour. Error bars indicate mean +/- SEM. 3 replicates were included for each treatment.

**Supplemental Figure 2. pH measurements of bovine synovial fluid (BSF) treatments and hyaluronic acid-supplemented solutions**. A pH meter was used to measure the pH of 10% BSF, 20% BSF, and 50% BSF in PBS. Additionally, the hyaluronic acid supplemented solutions were also assessed.. Error bars indicate mean +/- SEM. 3 biological replicates were included for each solution.

**Supplemental Figure 3. Gentamicin-killed *S. aureus* retains the capacity to aggregate in bovine synovial fluid (BSF).** *S. aureus* strain AH1726 was treated with gentamicin for 3-hours before the addition of 10% BSF. Time Lapse imaging was conducted to confirm aggregation occurred over time upon contact with BSF (**Fig. S3a**). After 1-hour of exposure, the average aggregate size was quantified using FIJI image analysis software for both gentamicin-treated and untreated bacteria (**Fig. S3b**). The bacterial killing was confirmed by CFU counts (**Fig. S3c**). Data points represent 3 replicates of gentamicin-treated bacteria and 2 replicates of untreated bacteria with 5 representative images collected for each replicate. Error bars indicate mean +/- SEM. Statistical significance was determined by Student’s T-test to compare the mean aggregate size between treated and untreated bacteria. ** p ≤ 0.01, ; ****p ≤ 0.0001

